# Tyrosinase from mushroom *Agaricus bisporus* as an inhibitor of the Hepatitis C virus

**DOI:** 10.1101/2020.12.23.424187

**Authors:** David Lopez-Tejedor, Rafael Clavería-Gimeno, Adrian Velazquez-Campoy, Olga Abian, Jose M. Palomo

## Abstract

Tyrosinases from both a commercial semi-purified *Agaricus bisporus* protein extract and directly isolated from white mushroom have been demonstrated to show antiviral activity against the Hepatitis C virus for the first time. The well-known tyrosinase from *A. bisporus (TyrAB)* of 45kDa and a newly discovered 50-kDa isoform from this tyrosinase *(Tyr50kDa)* have been tested. Cell toxicity and antiviral activity of tyrosinases in cultured Huh 5-2 liver tumor cells transfected with a replicon system (a plasmid that includes all non-structural Hepatitis C virus proteins and replicates autonomously) was determined. Native *TyrAB* was able to inhibit the replication of the hepatitis C virus without inducing toxicity in liver cells. In addition, the post-translational isoform of *Tyr50kDa* showed higher antiviral capacity than the former (up to 10 times greater), also exhibiting 10 times higher activity than the commercial drug Ribavirin^®^. This antiviral activity was directly proportional to the enzymatic activity of tyrosinases, since no antiviral capacity was observed for the inactive enzymes. The tyrosinases approach could represent a new antiviral inhibition mechanism, through a catalytic mechanism of selective hydroxylation of key role tyrosine residues in viral proteases. The tyrosinases directly extracted from fresh mushrooms (containing both tyrosinases) showed similar antiviral activity and, therefore, might provide low-cost drugs for the treatment of hepatitis C.

## Introduction

Infection by the hepatitis C virus (HCV) is a main health problem worldwide.^1^ HCV contains a small, single-stranded positive RNA that encodes for several proteins that are translated as a large polyprotein, which is processed in the endoplasmic reticulum of the infected cell by viral and host proteases in order to render functional the individual structural and non-structural viral proteins.^2^ Among these, the non-structural proteins^3–4^ NS2, NS3/4A, NS4B, NS5A and NS5B present great importance, because most of the drugs currently marketed with therapeutic indication (Figure 1) act through inhibition mechanisms directed towards one of these proteins.^5–9^

**Figure 1.**
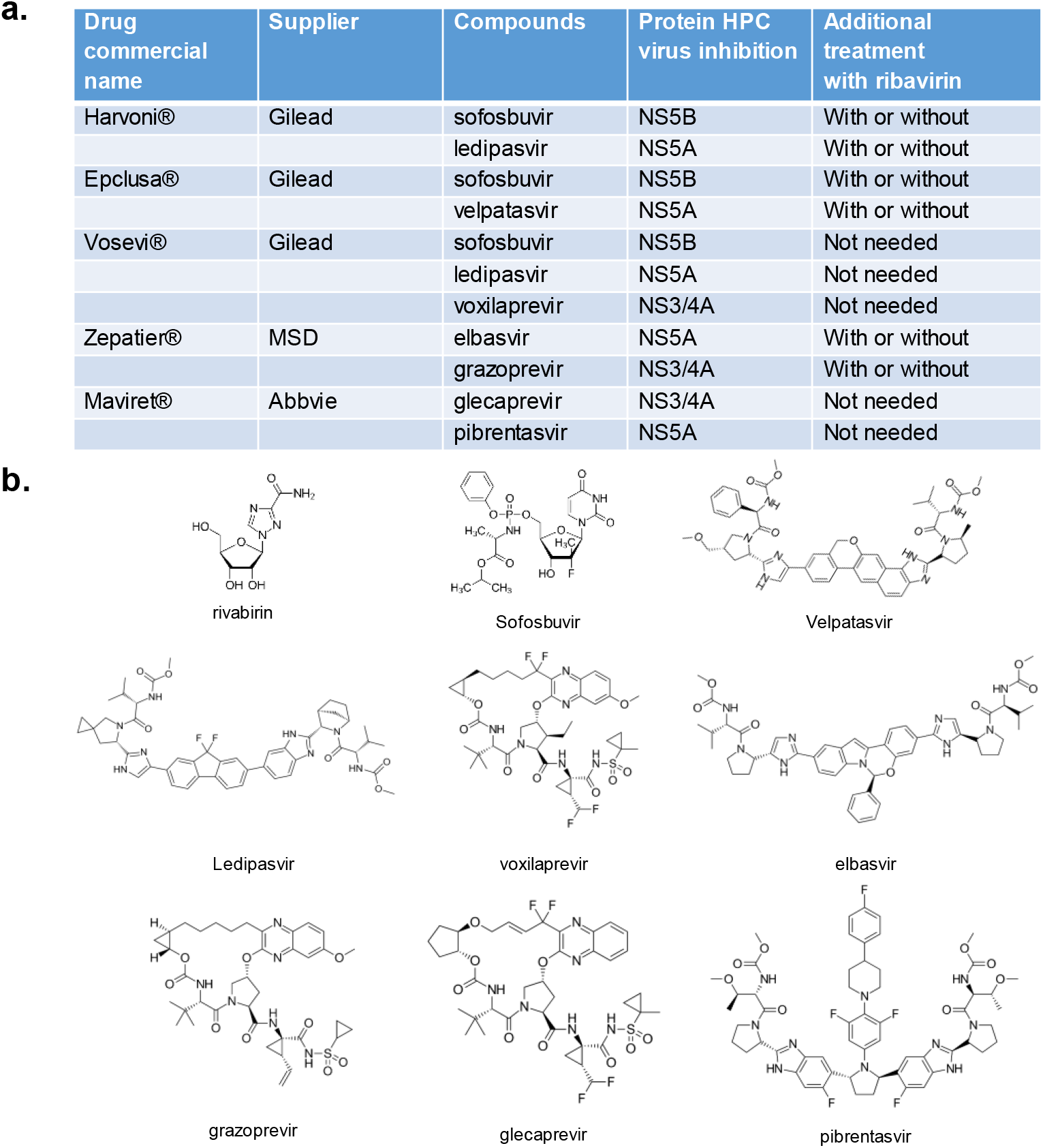
Currently marketed drugs against Hepatitis C virus. A) Pharmacological preparations. B) Chemical structures of active molecules.

NS2 is for assembly and replication due to its function as an autocatalytic cysteine protease.^10^ The *N*-terminal domain of the multifunctional NS3/4A (or just NS3) is the second viral protease responsible for processing most of the viral polypeptide, while the C-terminal domain of NS3 possesses a helicase/dNTPase function.^11^ NS4A is a small hydrophobic protein that serves as a cofactor for enhancing the activity of the NS3 serine protease.^12^ NS4B and NS5A are proteins involved in replication and assembly,^13^ and NS5B is the viral RNA-dependent RNA polymerase that forms a replication complex together with all other non-structural proteins.^14^

Currently, some of the best-known systems in the formulation of drugs against HCV consist of a mixture of several molecules (Figure 1).^15^ For example, Harvoni^®^, the last drug registered by Gilead in 2014, is a combination of sofosbuvir and ledipasvir, inhibitory compounds against NS5B and NS5A proteins, respectively (Table 1). Other commercial drugs consisting of a combination of compounds are: Epclusa^®^ (Gilead), Vosevi^®^ (Gilead), Zepatier^®^ (MSD), and Maviret^®^ (Abbvie) (Figure 1).^15^

**Table 1.** EC50 and LC50 of the different isoforms of tyrosinase purified from the commercial extract of *A. bisporus.*

|  | EC50<br>(µg/mL) | LC50<br>(µg/mL) |
| --- | --- | --- |
| *TyrAB* | 6-12 | ND (>>25) |
| *Tyr50kDa* | 1.2-2.5 | ND (>>25) |
| *Tyr50kDa* (37°C) | 10 | ND (>>25) |
| *Tyr50kDa* (80°C) inactive | ND (>20) | ND (>>25) |
| Rivabirin | 10 | ND (>>20) |

ND: not determined at the concentrations tested

There are chemical compounds, natural and synthetic, with antiviral capacity against HCV, for example, sesquiterpenoids extracted from the fermentation of the *Trichoderma harzianum* fungus^16^, peptide compounds^17^, GSK3082 analogues^18^ or aromatic drugs mimetics^19^.

However, currently available treatments for the hepatitis C virus are too expensive and many patients, especially from less developed countries, do not have access to them. They are treatments that cost around € 60,000,^20^ thus, even in developed countries, health budgets are exceeded, forcing governments to establish restrictive criteria for the prescription of these drugs. With a gross estimation of 150-200 million people infected by HCV worldwide, necessary and urgent to develop effective and alternative treatments to the existing ones that are also inexpensive.

In this work, we report for the first time a tyrosinase and one isoform present in the white mushroom *(Agaricus bisporus)* showing high inhibition capability against HCV. This presents an HCV inhibition mechanism different from those currently described (allosteric or orthosteric inhibition of non-structural HCV proteins).

## Results and Discussion

First, both toxicity and antiviral activity of the commercial *Agaricus bisporus* tyrosinase *(TyrAB)* extract against HCV in Huh-5-2 cells were evaluated. Tyrosinase is a heterotetramer (in fact, a dimer of heterodimers) made up of two H1L1H2L2 subunits, with two 45 kDa H subunits and two 12 kDa L subunits (Figure 2A).^21^ This extract was semipurified presenting only the H (active) subunits (H1H2 dimer) (Figure 2B).

**Figure 2.**
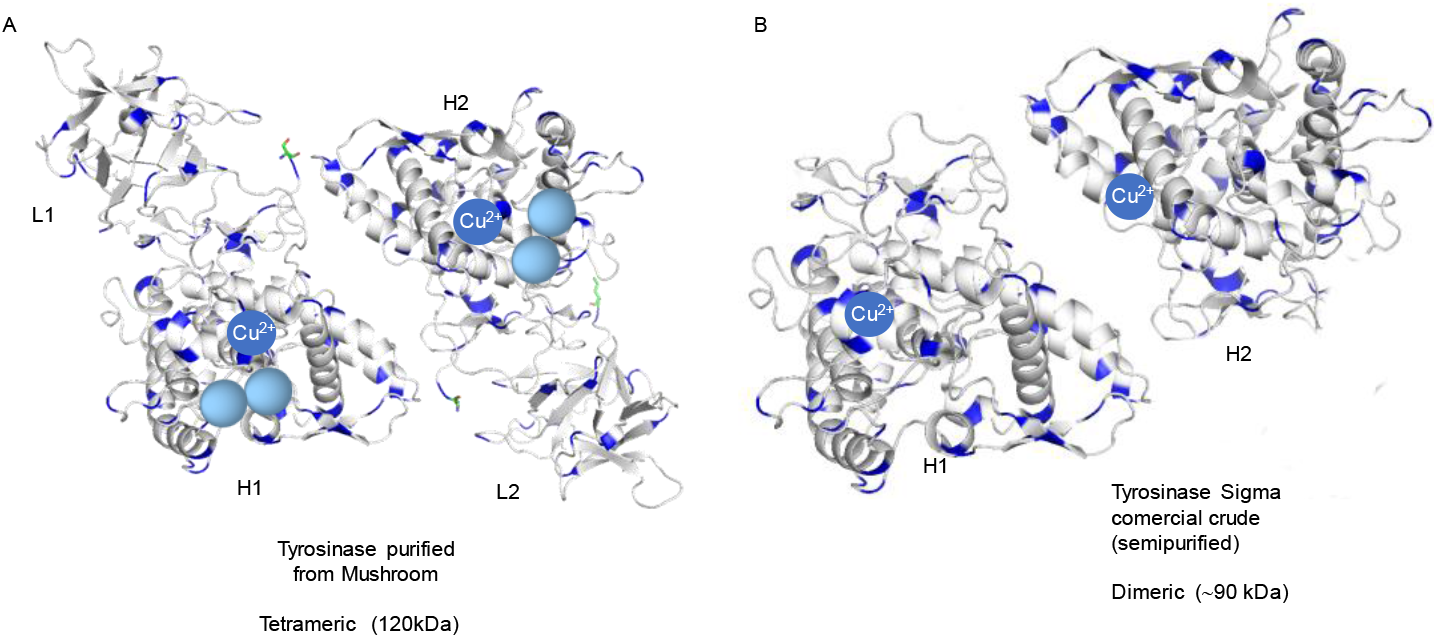
Structure of *Agaricus bisporus* tyrosinase

Furthermore, this semi-purified preparation contained an additional 50 kDa protein with tyrosinase activity *(Tyr50kDa)*, as identified by activity assay through native electrophoresis.^22^

The semi-purified Sigma extract (Tyr) was analyzed at two pH values (5 and 7), obtaining in both cases similar results, the extract did not present substantial toxicity towards liver cells and exhibits an unexpected inhibition activity of virus replication (EC50 of 20 μg/mL) (Figure 3).

**Figure 3.**
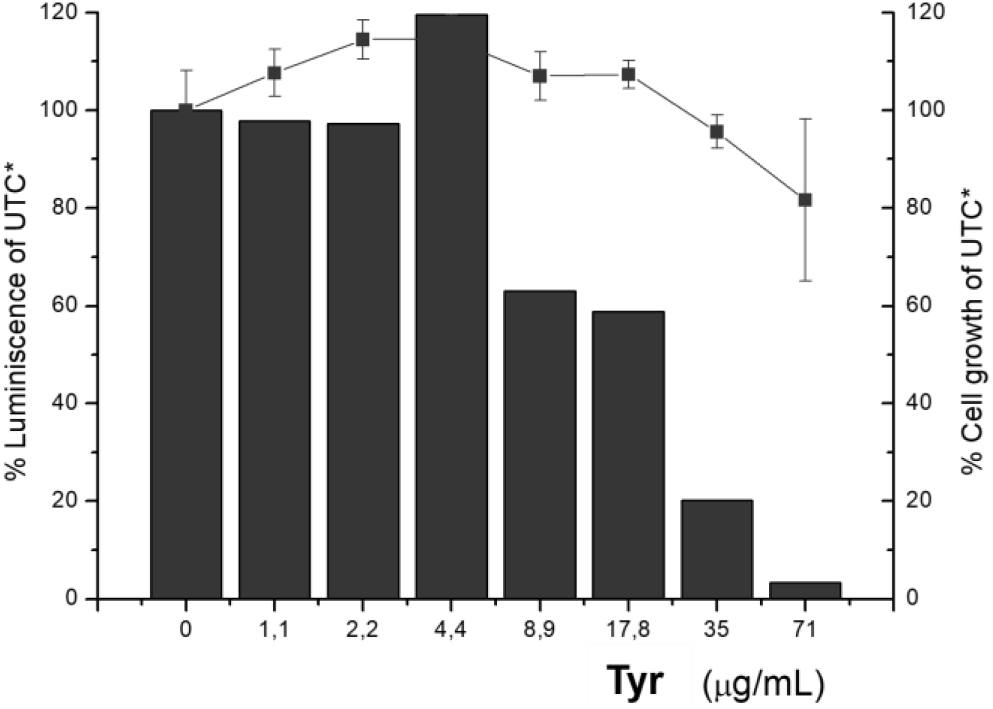
Inhibition of the Hepatitis C virus (HCV) replicon in cell assays. Evaluation of the potency and cytotoxicity of the protein in cell (Huh-5-2) assays. HCV replicon replication rate (luminescence, black bars) and cell survival (closed squares) were independently measured in cell culture by increasing protein concentration to determine EC50. UTC: untreated controls (no tyrosinase added).

To evaluate whether this activity specifically depends on the presence of tyrosinase, enzymes from the commercial extract was purified. Recently, we have reported the presence of a 50 kDa tyrosinase isoform *(Tyr50kDa)* that is more active than *TyrAB.^22^* Therefore, we proceeded to a semi-purification to separate both tyrosinases from the rest of the proteins from the extract and evaluate their antiviral activity.

This first preliminary purification process required the use of a detergent (Triton X-100). We observed that Triton X-100 concentrations higher than 0.25% (v/v) were toxic to the cells (Figure 4A). However, the purified samples containing less than 0.05% detergent, with both tyrosinases present, gave rise to an even better outcome: slight cellular toxicity and an EC50 for antiviral inhibition of 1 μg/mL (Figure 4B). This demonstrated that the active tyrosinases are responsible for the antiviral activity of the protein sample.

**Figure 4.**
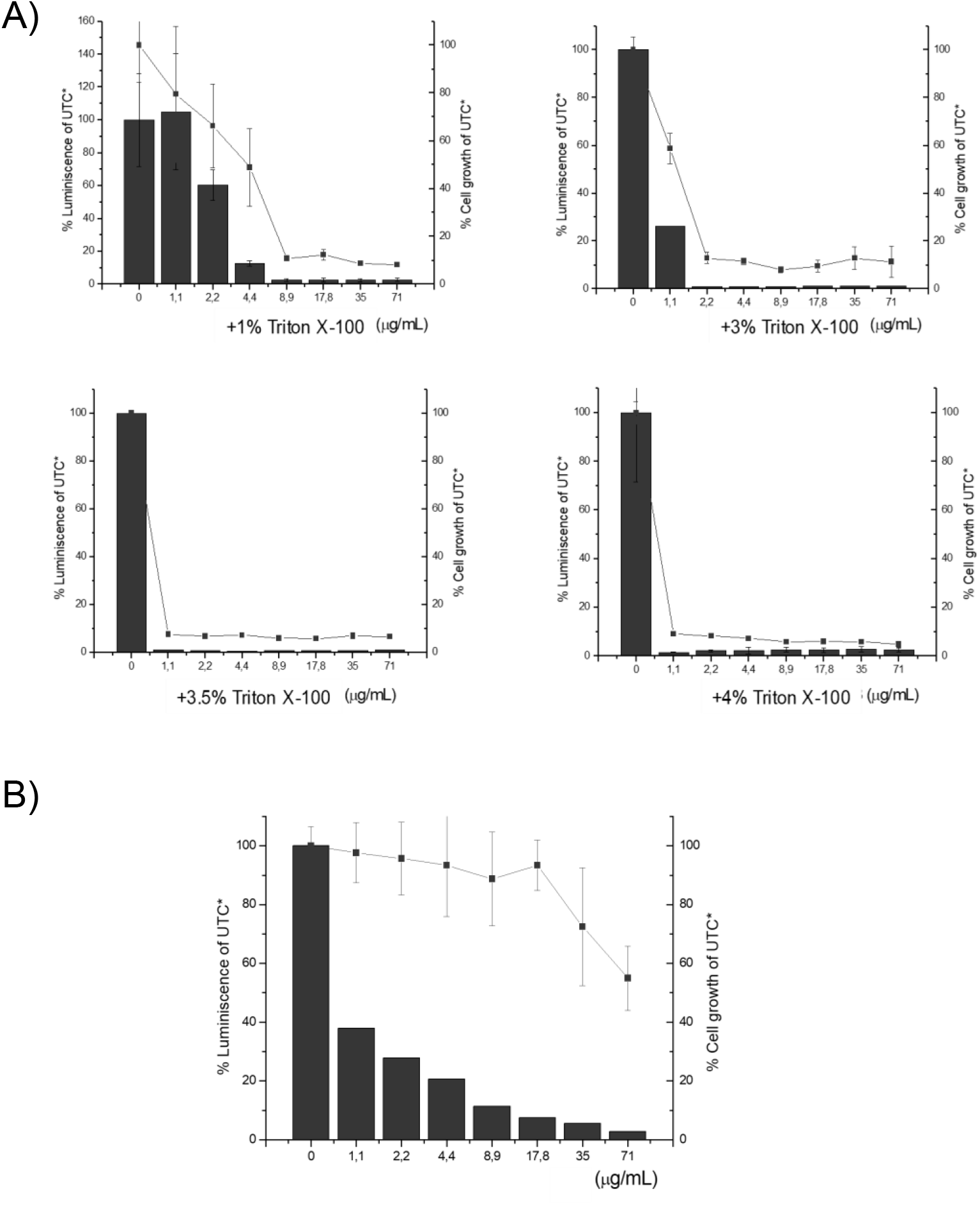
Effect of the presence of on the inhibition of the Hepatitis C virus (HCV) replicon in cell assays in the presence of Triton X-100. Evaluation of the potency and cytotoxicity of the protein in cell (Huh-5-2) assays. HCV replicon replication rate (luminescence, black bars) and cell survival (closed squares) were independently measured in cell culture by increasing protein concentration to determine EC50. A) 1-4% Triton X-100. B) The purified protein sample contained less than 0.05% TritonX-100.

To further confirm these results and to assess which tyrosinase variant was mainly responsible for the antiviral activity, both enzymes *(TyrAB* and *Tyr50kDa)* present in the extract were isolated as pure form using the recently described protocol.^22^ No cellular toxicity was observed for any of the enzymes, and both enzymes showed antiviral activity, although the *Tyr50kDa* isoform was responsible for most of the activity, with an inhibition capability of up to 10 times compared to that observed for *TyrAB* (EC50 of 1.22.5 μg/mL for *Tyr50kDa* compared to 6-12 μg/mL for *TyrAB)* (Figure 5, Table 1).

**Figure 5.**
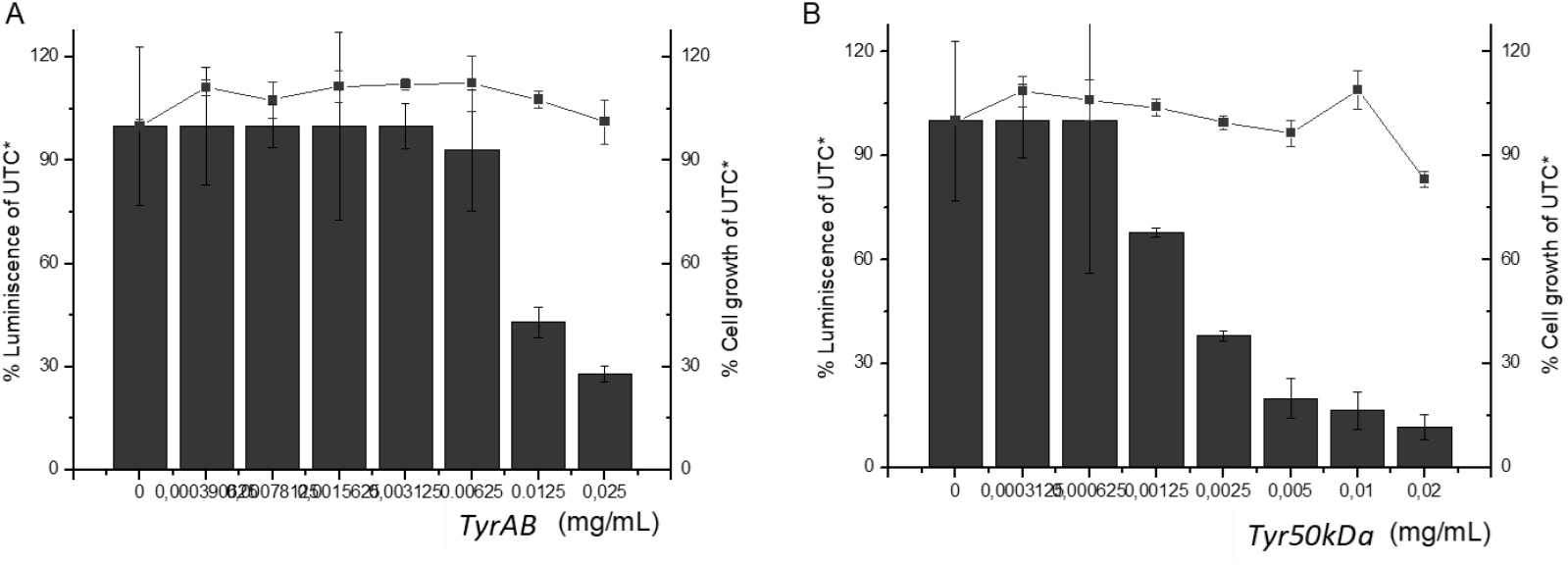
Inhibition of the Hepatitis C virus (HCV) replicon in cell assays. Evaluation of the potency and cytotoxicity of pure isoforms in cell (Huh-5-2) assays. HCV replicon replication rate (luminescence, black bars) and cell survival (closed squares) were independently measured in cell culture by increasing protein concentration to determine EC50 for (A) purified TyrAB and (B) purified Tyr50kDa.

This result indicates that tyrosinase from mushroom exhibits inhibitory activity against HCV, but also reveals that the isolated isoform (*Tyr50kDa*) recently discovered in our laboratory was responsible for most of the antiviral activity. Analysis and sequencing of the electrophoresis band corresponding to the pure *Tyr50kDa* suggested it corresponds to a post-translationally modified isoform of *TyrAB,* in particular, glycosylation,^22^ since the protein sequence exactly matches that of *TyrAB.*

These results provide evidence for a first time reported mechanism for inhibiting viral replication based on an enzymatic activity, in this case, based on the selective oxidation of tyrosine to L-DOPA and dopaquinone (Figure 6).

**Figure 6.**
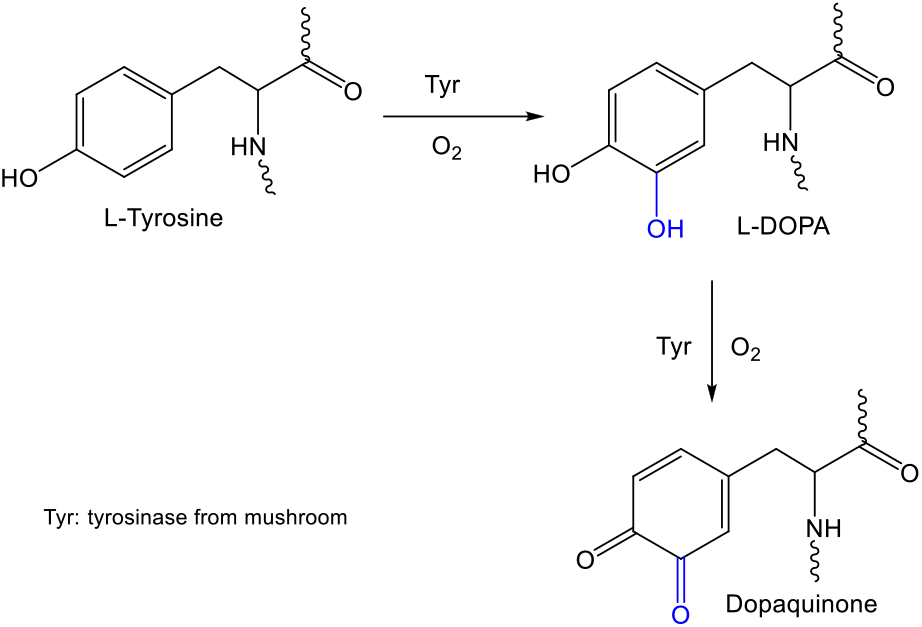
Selective oxidation of tyrosine residues catalyzed by tyrosinase from mushroom *A. bisporus.*

To further verify that this new mechanism of action for inhibition of HCV replication is specifically dependent on tyrosinase activity, the inhibitory effect of partially and totally inactivated tyrosinase was evaluated (Figure 7). Purified *Tyr50kDa* was incubated at 37°C or at 80°C for 24 h. The catalytic activity involving L-DOPA as a substrate showed that incubation at 37°C led to 80% reduction of activity, while incubation at 80°C yielded a completely inactive enzyme. This was additionally confirmed by assessing the tertiary structure loss in the enzyme upon temperature treatment by circular dichroism. Tyrosinase incubated at 37°C gave an EC50 value of 10 μg/mL (10-fold larger than that of the fully active enzyme), while that inactivated at 80°C did not inhibit viral replication at all (Table 1). Therefore, the inhibitory mechanism of action of tyrosinase relies on its enzymatic activity.

**Figure 7.**
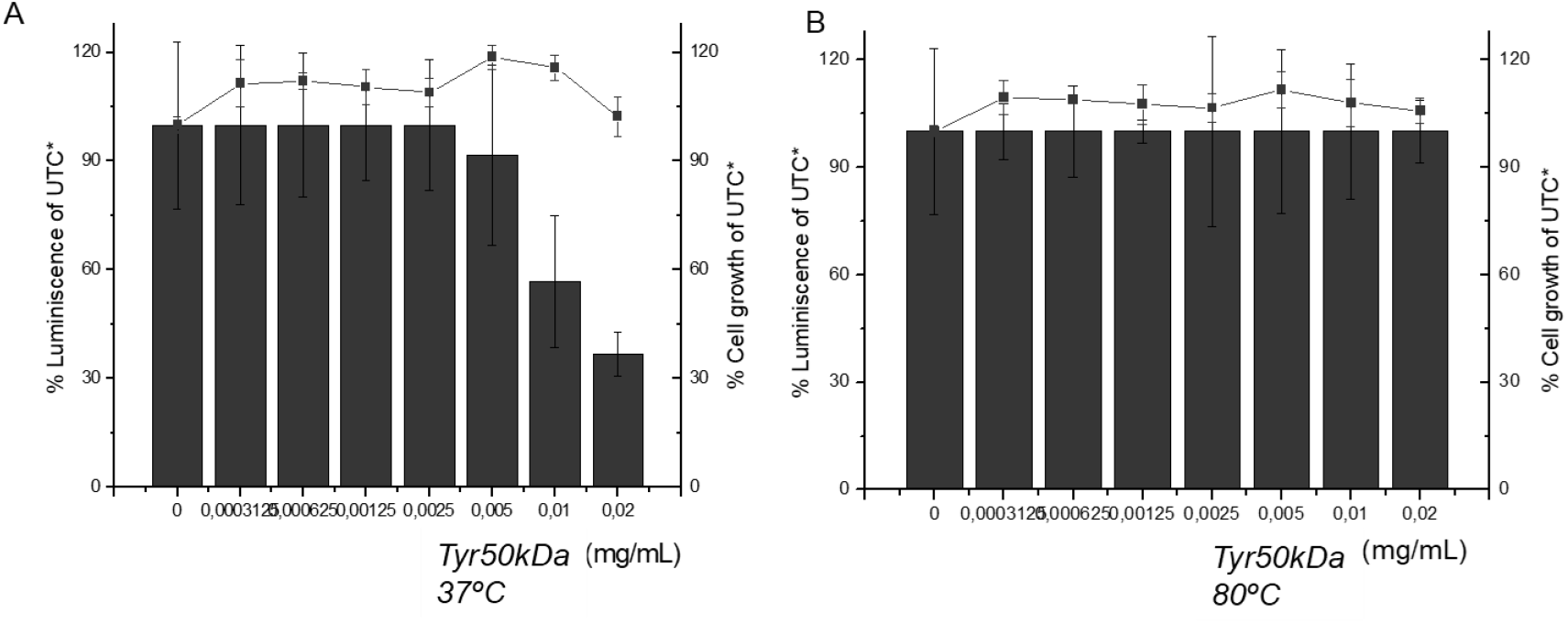
Inhibition of the Hepatitis C virus (HCV) replicon in cell assays. Evaluation of the potency and cytotoxicity of temperature-inactivated Tyr50kDa in cell (Huh-5-2-) assays for: A) partially inactivated at 37°C, and (B) completely inactivated at 80°C *Tyr50kDa*. HCV replicon replication rate (luminescence, black bars) and cell survival (closed squares) were independently measured in cell culture by increasing protein concentration to determine EC50.

According to these results, the mechanism of action of this protein sample is dependent on the selective oxidation of a surface tyrosine to L-DOPA and subsequent dopaquinone of proteins involved in the replication of the virus (NS3/4A, NS4B or NS5A/B).

From the structures of both NS3 and NS5A, some tyrosine residues as candidates for selective oxidation can be spotted on their surface. For example, the oxidation of Tyr93 in NS5A^23^ (Figure 8A) could produce the imminent reaction with the amino terminal of the protein, thus precluding its location in the membrane and, therefore, blocking the replication of the virus. Also, the alteration of the catalytic activity of NS3 or its structural change (Figure 8B) affecting its interaction with NS4A, could be instrumental for blocking the replication of the virus. These hypotheses must be proved in future work.

**Figure 8.**
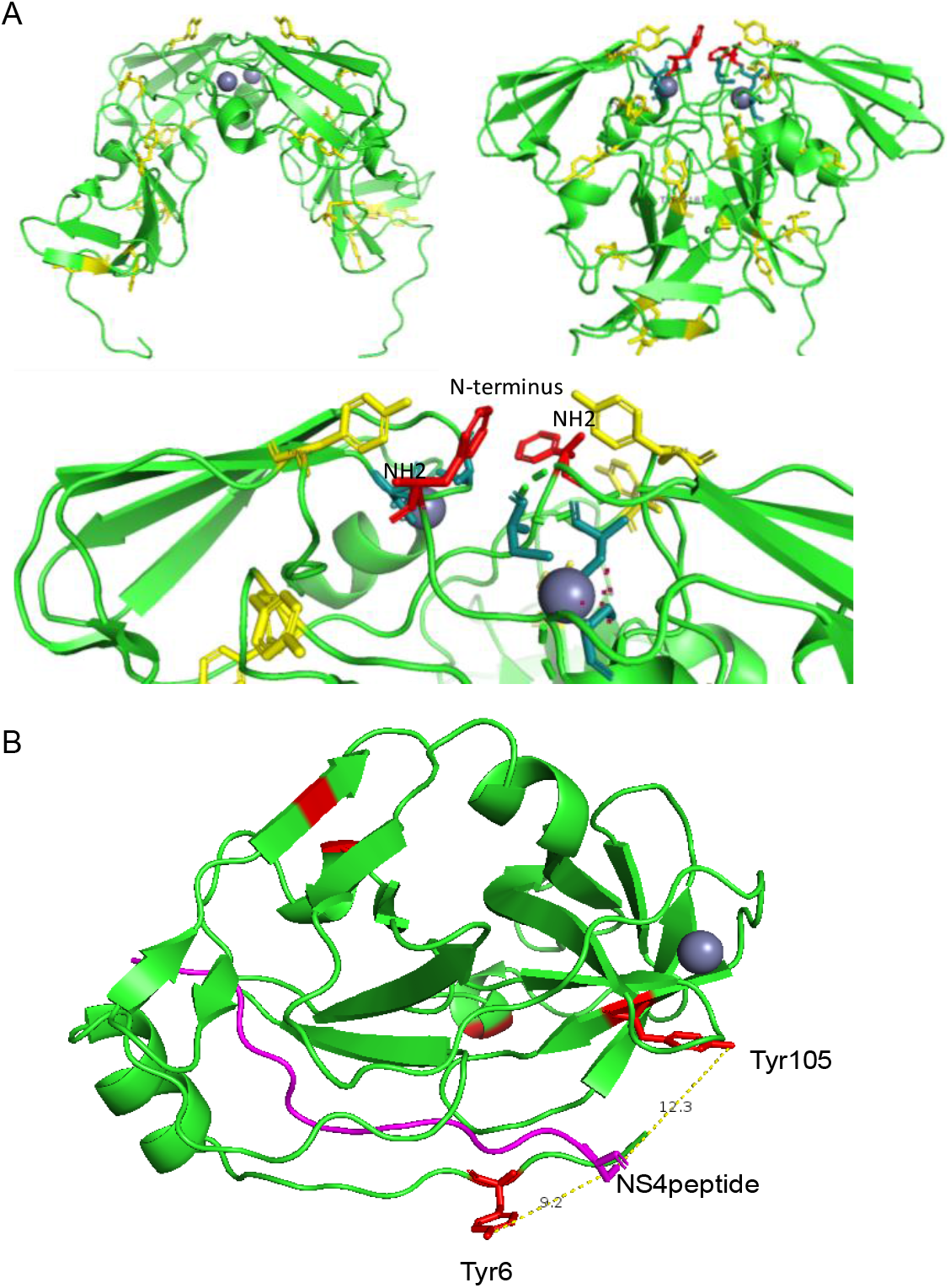
(A) 3D structure of the phosphoprotein protein NS5A, in dimeric form. Seven tyrosine residues (yellow) in each monomer, with one tyrosine in the N-terminal end (red). (B) 3D structure of the NS3 protease including the NS4 peptide (pink), showing the tyrosine residues in red.

As a key comparison, under the same conditions, ribavirin (a drug still used in combination with other drugs, such as Harvoni^®^ in the treatment of patients with cirrhosis) presented an EC50 of 10 μg/mL. This means that *Tyr50kDa* was almost 10 times more potent than ribavirin (Table 1).

In addition, the antiviral activity of a protein extract produced directly from the mushroom (*A. bisporus*) was evaluated. By means of a simple extraction, together with solubilisation, precipitation, and dialysis steps, it was possible to obtain a purified protein sample of the two enzymes. In this case, the extraction and semi-purification method made it possible to obtain tyrosinase in its tetrameric form with all its subunits; thus, effect of the presence or absence of the L subunits of the heterodimer could be assessed. This semi-purified extract did not present cellular toxicity, giving a value for inhibition of viral replication similar to that obtained previously for the commercial extract, again corroborating its greater inhibitory capability of the virus in comparison with ribavirin (Table 2).

**Table 2.** EC50 and LC50 of the extract purified directly from mushroom and containing both tyrosinases (Tyr: *TyrAB* + *Tyr50kDa*).

|  | **EC50**<br>**($\mu\text{g/mL}$)** | **LC50**<br>**($\mu\text{g/mL}$)** |
| --- | --- | --- |
| Ribavirin | 10 | ND (>>20) |
| Tyr (protein extracted from fresh mushroom) | 2.5 | 10 |

## Conclusion

We have shown that two tyrosinases (a well-known enzyme plus a new 50 kDa isoform) present in a mushroom extract (*A. bisporus*) show antiviral activity against the Hepatitis C virus. The inhibitory activity relies on the catalytic mechanism for selective hydroxylation of tyrosine. The recently identified isoform, *Tyr50kDa*, is more active than the known isoform, and almost 10 times more active than the drug Ribavirin.

In addition, it has been shown the antiviral activity of both proteins can be detected in a similar way in a commercial semi-purified extract and in an extract directly produced from fresh mushrooms.

This protein preparation might become a promising therapeutic protein, which could be used as a substitute for some drugs or used in combination with others, such as Sofosbuvir in Harvoni^®^. It would be much more economic due to its natural origin, which could reduce the production costs. Also, this protein could be employed for covering surfaces with antiviral particles for prophylactic/hygienization purposes.

This new viral inhibition mechanism of action discovered for tyrosinase could also be effective for other virus, e.g., other hepatitis viruses, Dengue or Zika virus, thus, being a promising broad spectrum pharmacological agent.

## Experimental part

### Purification of the different tyrosinases from a semi-purified commercial extract of *Agaricus bisporus* from Sigma

The purification protocol has been optimized to purify two different active tyrosinase isoforms in dimeric form (H subunits) (45 kDa, 50kDa) *(TyrAB, Tyr-50kDa)* from commercial raw Sigma-Aldrich^®^ powder. Briefly, 1 mg of commercial lyophilized tyrosinase was dissolved in 4 mL of distilled water. 0.5 g of octadecyl-Sepabeads^®^ (C18-1) was added to this solution and the mixture was incubated for 2 h. After that, the mixture was vacuum filtered, the supernatant was recovered and sodium phosphate buffer was added at pH 7 to a final concentration of 100 mM (for example, 0.5 ml of 2M phosphate buffer solution at pH 7 Triton ™ X-100 was subsequently added to a final concentration of 0.07% (w / v). 0.5 g of fresh octadecyl-Sepabeads^®^ (C18-2) was added to this supernatant and the mixture was incubated for 3 h. After this time, the supernatant contained only the *Tyr-50kDa* enzyme, while *TyrAB* and a slight amount of *Tyr-50kDa* were on the solid (confirmed by SDS-PAGE). A previous semi-purification was also carried out by desorption of the C18-2 derivative using different concentrations of Triton X-100. Finally, the soluble protein samples were purified to reduce the amount of detergent in the sample by ultrafiltration with Amicon Ultra 10kDa (<0.05%).

### Tyrosinase extraction procedure directly from the mushroom

25 g of white mushrooms (*A. bisporus*) (from Eroski supermarkets) were cut into small pieces and added to 50 ml of cold acetone. The mixture was kept under paddle stirring for 30 minutes in an ice bath. Then, the mixture was centrifuged at 7000 rpm for 20 minutes and the pellet was recovered, discarding the solution. This solid was suspended in 25 ml of distilled water and the mixture was incubated for 1 hour and then centrifuged at 8000 rpm for 40 min. After filtration, the saline precipitation process was performed adding 60% (w/v) of ammonium sulfate, adding it little by little and then 1 h of incubation. After that, the mixture was centrifuged at 8000 rpm for 40 minutes and the solid was recovered and stored at −20 ° C. Solid was dissolved in water and dialysis was performed before used.

### *In vitro* L-Dopa activity assay

Tyrosinase activity was tested in the presence of 2 mL of 1 mM L-DOPA in 0.1 M phosphate buffer pH 7 at room temperature using a V-730 spectrophotometer (Jasco), measuring the increase in absorbance of aminochromes at 475 nm caused by 40 μL of enzyme solution, and taking the initial speed, between 10 and 70 seconds of the reaction. One unit of enzyme activity (U) was defined as the amount of enzyme that causes a 0.001 increase in absorbance /min at 25 ° C.

### Evaluation of cellular toxicity and antiviral activity

Huh-5-2 (Lunet, liver tumor cells) cells were transfected with a plasmid containing a replicon system (pseudovirus including all the non-structural proteins from the Hepatitis C virus): NS2, NS3, NS4A, NS4B, NS5A, and NS5B). Because the replicon system does not contain structural proteins, it is not infective; however, it replicates autonomously and allows quantifying, through a reporter gene encoding for for the enzyme luciferase, the rate of replication within the cells as proportional to the luminescence signal after ading an appropriate luciferase substrate. In the case of inhibition of replication, a reduction of the luminescence signal will be observed compared to the corresponding control.

The Huh5-2 cell culture was performed using DMEM with phenol red as the culture medium, supplemented with Fetal Bovine Serum (FBS), penicillin, glutamine, streptavidin, non-essential amino acid, and 500 μg/mL geneticin (G418) from Invitrogen (selection of cells with a replicon system inside them). The culture conditions were: temperature (37°C), 5% CO2, 95% air and 1 atm of pressure.

To determine both the antiviral activity of tyrosinases and their cellular toxicity, 96-well plates were seeded with 7000 cells/well (using DMEM without phenol red) with 100 μL final volume, in which increasing serial concentrations of tyrosinases (from 0 to 0.2 mg/mL) were tested in triplicate for each of the concentrations. The cells were incubated for 72 h with tyrosinases, after which both their cytotoxicity and their antiviral activity were evaluated (in duplicate plates).

For cytotoxicity, in one of the plates, the cellular metabolism level of each of the wells was determined using the Cell Titer 96^®^AQueous Reagent (Promega) in an absorbance plate reader (Synergy HT Multi-Modal Microplate Reader with Gen5 Data Analysis software, BioTek). The supernatant from the wells was removed, and 20 μL of Cell Titer 96^®^AQueous were diluted 1:4 in the same DMEM culture medium, but with no RF added, and incubated for 3 h at 37°C. The absorbance was measured at 490nm and 800nm, and the difference between these absorbance values reports the substrate metabolized by viable cells in the culture. Each tyrosinase concentration was evaluated in triplicate, and the values obtained were averaged. The relationship between the tyrosinase concentration and the percentage of absorbance were plotted (taking as 100% those values in the control wells with no tyrosinase treatment). The LC50 was calculated as the tyrosinase concentration at which cell viability was reduced by 50% (with respect to that obtained in the absence of tyrosinase treatment).

For antiviral activity, the luminescence of each of the wells was determined using the Bright-Glo™ Luciferase Assay System Reagent (Promega) in a luminescence plate reader (Synergy HT Multi-Modal Microplate Reader with Gen5 Data Analysis software, BioTek). Without removing the supernatant, 30 μL of reagent was added to each well, and the luminescence signal was measured. The luminescence signal was proportional to the amount transcribed reporter gene. Each tyrosinase concentration was evaluated in triplicate, and the values obtained were averaged. The relationship between the concentration of tyrosinase and the percentage of luminescence was plotted graph (taking as 100% those values in the control wells with no tyrosinase treatment). EC50 was calculated as the tyrosinase concentration at which viral replication is reduced by 50% (with respect to that obtained in the absence of tyrosinase treatment).

## Acknowledgments

This work has been sponsored by the Spanish National Research Council (CSIC) (project 201880E011), Spanish Ministry of Economy and Competitiveness and European ERDF Funds (MCIU/AEI/FEDER, EU) (BFU2013-47064-P and BFU2016-78232-P to A.V.C.); Miguel Servet Program from Instituto de Salud Carlos III (CPII13/00017 to O.A.); Fondo de Investigaciones Sanitarias from Instituto de Salud Carlos III and European Union (ERDF/ESF, “Investing in your future”) (PI15/00663 and PI18/00349 to O.A.); Diputación General de Aragón (Protein Targets and Bioactive Compounds Group E45_17R to A.V.C. and Digestive Pathology Group B25_17R to O.A.); and Centro de Investigación Biomédica en Red en Enfermedades Hepáticas y Digestivas (CIBERehd).

